# Methylation Array Signals are Predictive of Chronological Age Without Bisulfite Conversion

**DOI:** 10.1101/2023.12.20.572465

**Authors:** Hunter L. Porter, Victor A. Ansere, Ram Babu Undi, Walker Hoolehan, Cory B. Giles, Chase A. Brown, David Stanford, Mark M. Huycke, Willard M. Freeman, Jonathan D. Wren

## Abstract

DNA methylation data has been used to make “epigenetic clocks” which attempt to measure chronological and biological aging. These models rely on data derived from bisulfite-based measurements, which exploit a semi-selective deamination and a genomic reference to determine methylation states. Here, we demonstrate how another hallmark of aging, genomic instability, influences methylation measurements in both bisulfite sequencing and methylation arrays. We found that non-methylation factors lead to “pseudomethylation” signals that are both confounding of epigenetic clocks and uniquely age predictive. Quantifying these covariates in aging studies will be critical to building better clocks and designing appropriate studies of epigenetic aging.

## I. Introduction

Epigenetic alterations and genomic instability are each hallmarks of the aging process [1]. Prior studies have shown that high-throughput epigenetic DNA modification assays, such as DNA methylation arrays, can be used to make “epigenetic clock” models that are predictive of chronological age[2, 3]. Further studies on DNA methylation in aging led to the development of second-generation clocks purporting to measure “biological aging” [4, 5]. The relationship between predicted “accelerated aging” and age-related disease and dysfunction is unclear, and often contradictory [6, 7]. Nonetheless, clock studies suggest a molecular mechanism that might be a surrogate endpoint for evaluating anti-aging interventions in humans [8]. The use of epigenetic clocks is rapidly on the rise [4, 5, 9-12], and has even inspired new theories of aging[10]. Most methylation studies, however, rely on sodium bisulfite conversion to identify the methylation status of CpGs [13].

Sodium bisulfite catalyzes the deamination of cytosine to uracil[14]. This method assumes that thymines observed at what are cytosines in the reference genome (including microarray probes) are due to bisulfite conversion of unmodified cytosine (Figure 1A). A potential confound is that accumulation of mutations with aging is well known[15]. More specifically, there is significant evidence that deamination mutations accumulate with aging[16] and may even be age predictive[17]. As well, germline mutations can be predicted from alterations to methylation array signals[18], but the somatic mutation signal does not follow allelic ratios and may be difficult or impossible to identify within the same data. Mutations in CpG dinucleotides are predictive of time at the taxonomic scale [11], and the mutation rate appears to be related to the methylation level of cytosines [19]. Cytosine to thymine (C=>T) substitutions are the most abundant in the genome with aging [16, 20]. C=>T mutations were previously linked to patient age in a study analyzing mutations in human cancers; they found two “clocks” in mutational signatures and proposed an etiology of cytosine deaminations correlated with patient age [21].

**Figure 1.**
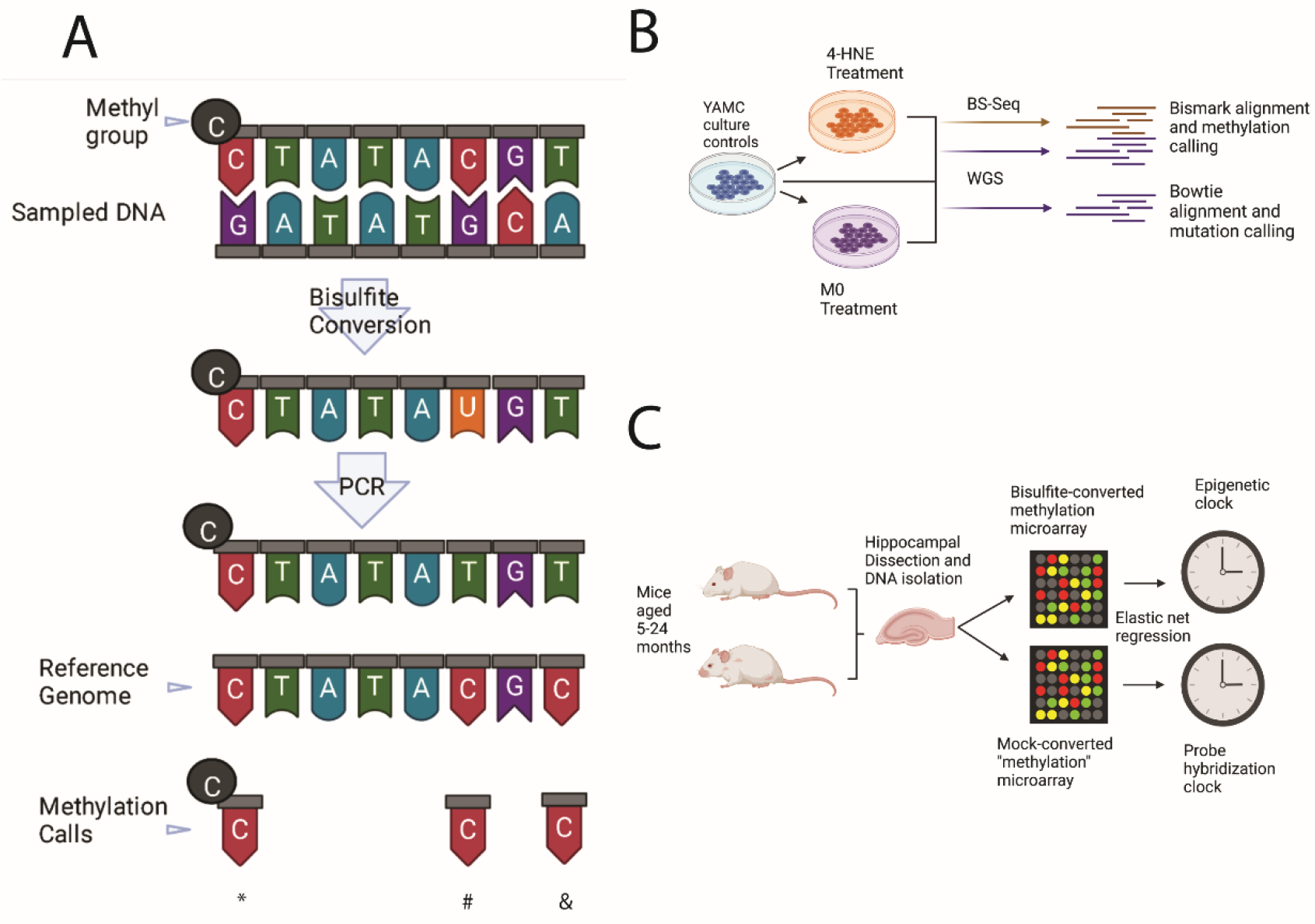
Conceptual framework and methodology for relating DNA methylation, somatic mutations, and age prediction models. **A)** Diagram showing how bisulfite treatment is used to make methylation calls. Standard methylation calls (denoted by *) and unmethylated calls (#) are contrasted with an example of a C=>T mutation being read as unmethylated (&) **B)** Schematic of generation of paired mutation and methylation profiles using Illumina sequencing. **C)** Schematic of generation of paired converted and unconverted “methylation” data using methylation microarrays and subsequent epigenetic clock analysis.

Thus, it should not be controversial to suggest part of the age-related signal from epigenetic clocks is due to mutations, but the degree and effects are unclear. If aging creates many somatic C=>T mutations, they might induce an apparent “genomic hypomethylation” with aging, which was previously a commonly accepted theory [22]. However, T => C mutations are almost as abundant in aged samples and may create new methylatable loci [16]. The array data, where epigenetic clocks are most thoroughly studied in humans, further complicate the equation by hybridizing physical DNA molecules – allowing even other mutations to influence affinity [23]. Answering this is key to understanding the biological ramifications of epigenetic clocks, and the behavior we might expect if they are to be a biomarker of aging that is responsive to treatment[24].

In our previous report [25], we identified that 1) epigenetic clocks can be trained to use a variety of redundant loci to predict age; 2) epigenetic clocks use loci that exhibit age-related changes much smaller than we expect to be physiologically impactful; and 3) the age-predictive loci are enriched outside of coding regions of the genome and are somewhat associated to annotated eQTLs. These observations, and the previously described context in the literature, led us to question if age-predictive methylation changes could be explained by accrual of mutations in regions with low selection pressure, such as in pseudogenes in taxonomic analyses[26].

To approach these questions, we first attempted to understand how somatic mutations relate to readouts of DNA methylation by paired whole genome sequencing (i.e., untreated, to detect mutations) and bisulfite-sequencing from mouse colon epithelial cell lines transformed into cancer cells by polarized macrophages (M φ) or 4-hydroxy-2-nonenol (4-HNE) (Figure 1B). **Our first hypothesis was that somatic mutations would be enriched in regions of altered DNA methylation, and, further, that mutations within unconverted sequencing data would be read as loss of methylation when analyzed using bisulfite analysis tools**. Next, we tested if microarrays could measure signals other than DNA methylation and produce age-predictive clocks. To this end, we generated Illumina Mouse Methylation array data using bisulfite converted DNA or DNA subjected to a mock conversion process lacking bisulfite (unconverted). These data were isolated from the hippocampi of mice between 5-24 months of age with a total N of 38. We then trained epigenetic clocks using converted and unconverted data (Figure 1C). **Our second hypothesis was that we could generate models with similar age predictions using paired converted and unconverted DNA signals from methylation arrays**.

## II. Results

### Mutations can be read as losses of DNA methylation in bisulfite sequencing data

To evaluate if mutations could affect methylation readouts relevant to aging biology and epigenetic clocks, we first sought to quantify their relationship in a simpler context. Using a cell line that had been transformed into cancer clones by induced mutagenesis, we expected an increase in the preponderance of mutations in the genome. High-coverage sequencing (WGSS >30X, BOCS > 15X, Joint > 10X) allowed us to assay 145,667 loci detected in both unconverted and converted sequencing data (Figure 2). Consistent with our expectation, we observed that WGSS processed through Bismark[27] identifies almost all CpG nucleotides as “methylated”, since they are cytosines in both the reference genome and collected sample (Figure 2A). Further, filtering to loci with independently called mutations revealed that the mutation-adjacent loci have a different methylation distribution (Figure 2B). Interestingly, the WGSS data showed a distribution comparable to real methylation near the mutated loci, congruent with our hypothesis that deamination mutations could produce apparent losses of methylation. We observed that the correlation between methylation signal and “pseudomethylation” from WGSS was low (pearson *r* = 0.0914) in the overall data (Figure 2C). The correlation is further depleted when removing any loci with mutations (pearson *r* = 0.0400), and greatly enriched in mutation-adjacent loci (pearson *r* = 0.5750) (Figure 2D). We then tried to control for the shift in methylation data induced by the mutations to assess how different the apparent methylation levels would be if we accounted for mutations. We rescaled the methylation values by dividing by apparent methylation by pseudomethylation within each sample at each position; this transformation makes loci with pseudomethylation equal to methylation into the new 100% methylated call. This transformation revealed that many loci were erroneously reported as losses of methylation when in reality they were genomic mutations (Figure 2E). At whole-genome scale, most of the loci are unaffected by mutation readouts (Figure 2F), but at some loci, mutations shift apparent methylation levels by 50+%. These results suggest that mutations do in fact influence methylation readouts, and it may be possible to deconvolve methylation signals with paired mutation distributions.

**Figure 2.**
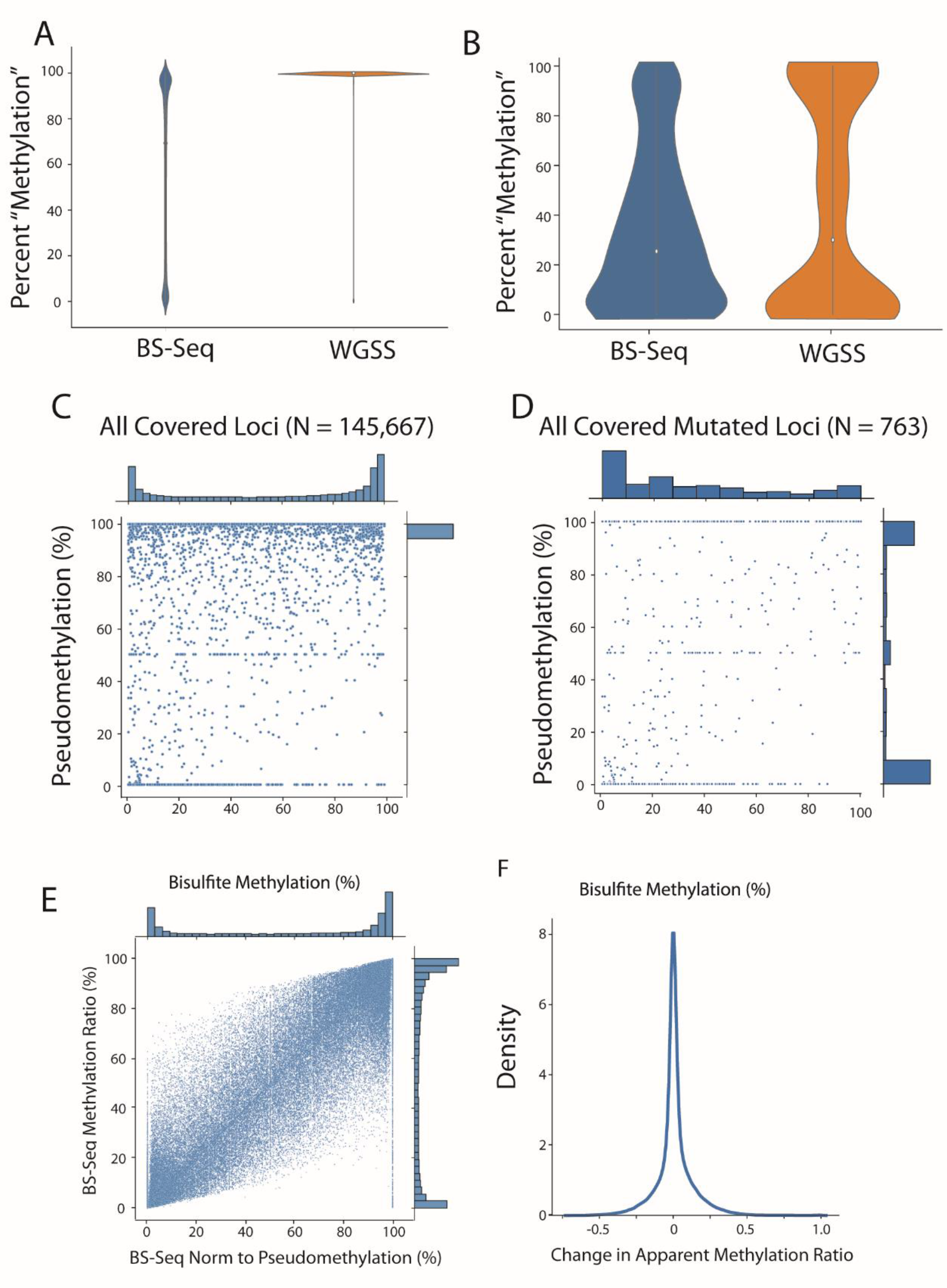
Somatic mutations in cancer transformation can be read as loss of methylation and produce methylation-like signals. Bisulfite Oligonucleotide Capture Sequencing (BOCS) and Whole-genome shotgun sequencing (WGSS) data were analyzed using Bowtie2 and Bismark. **A** Overall distribution of methylation (BS-Seq) and pseudomethylation (WGSS) at all sufficiently-covered loci. **B** Distribution of methylation and pseudomethylation in loci with confident mutation calls. **C** Relationship between pseudomethylation and methylation at all loci (pearson *r* = 0.0914). **D** Relationship between pseudomethylation and methylation at confident mutation-called loci (pearson *r* = 0.5750). **E** Relationship between standard bisulfite and bisulfite sequencing adjusted for pseudomethylation level at each locus. **F** Distribution of changes in apparent methylation (methylation – (methylation/pseudomethylation)) across all loci.

### Epigenetic clocks can be trained on microarray data without bisulfite conversion of DNA

Given our demonstration that mutations can affect methylation readouts from a mutation-rich context (cancer transformation) in sequencing data, we next interrogated how these signals may affect epigenetic clock predictions. While methylation arrays cannot measure mutations directly, we hypothesized that alterations to the sequence may occur during aging, other time-related processes[28], or during the extra-bisulfite processing steps, that could be detected by arrays and predictive of age. To this end, we generated two datasets over the same 38 samples. The first dataset was processed as indicated by Illumina [13], generating apparent methylation data from bisulfite-converted DNA samples (converted), while the second was generated using the same workflow but without addition of sodium bisulfite (unconverted). These datasets were then used to train and evaluate “epigenetic” clocks.

To ensure robustness of the observed age predictions, we iteratively split our data into 75/25 train/test splits, generating 100 epigenetic clocks for each converted and unconverted dataset(Figure 3). These models allowed us to interrogate the accuracy of age prediction from both datasets, and to ask questions about the loci selected by each type of aging model. We first observed comparable performance between converted (Figure 3A) and unconverted (Figure 3B) models trained on the same train/test split. Further, we interrogated the accuracy of the models in terms of R^2^ between predicted and chronological age (Figure 3C) and the Mean Absolute Error (MAE) of the predictions (Figure 3D). We observed an overall median difference in R^2^ of -0.08325 and median difference in MAE of 0.2748 months. These differences are well within one standard deviation of the R^2^ values (0.209) and the MAE (0.667). Interestingly, the unconverted clocks tended to select fewer loci (mean non-zero coefficients = 392) than converted clocks (mean non-zero coefficients = 1,289) despite them having comparable training set sizes (17,146 unconverted loci, 13,138 converted loci). Taken together, these results show we can generate “epigenetic” clocks even without the application of sodium bisulfite which encodes the methylation status into a discernable base difference. However, it is still undetermined if the methylation array signal detected with unconverted DNA is directly caused by DNA mutations.

**Figure 3.**
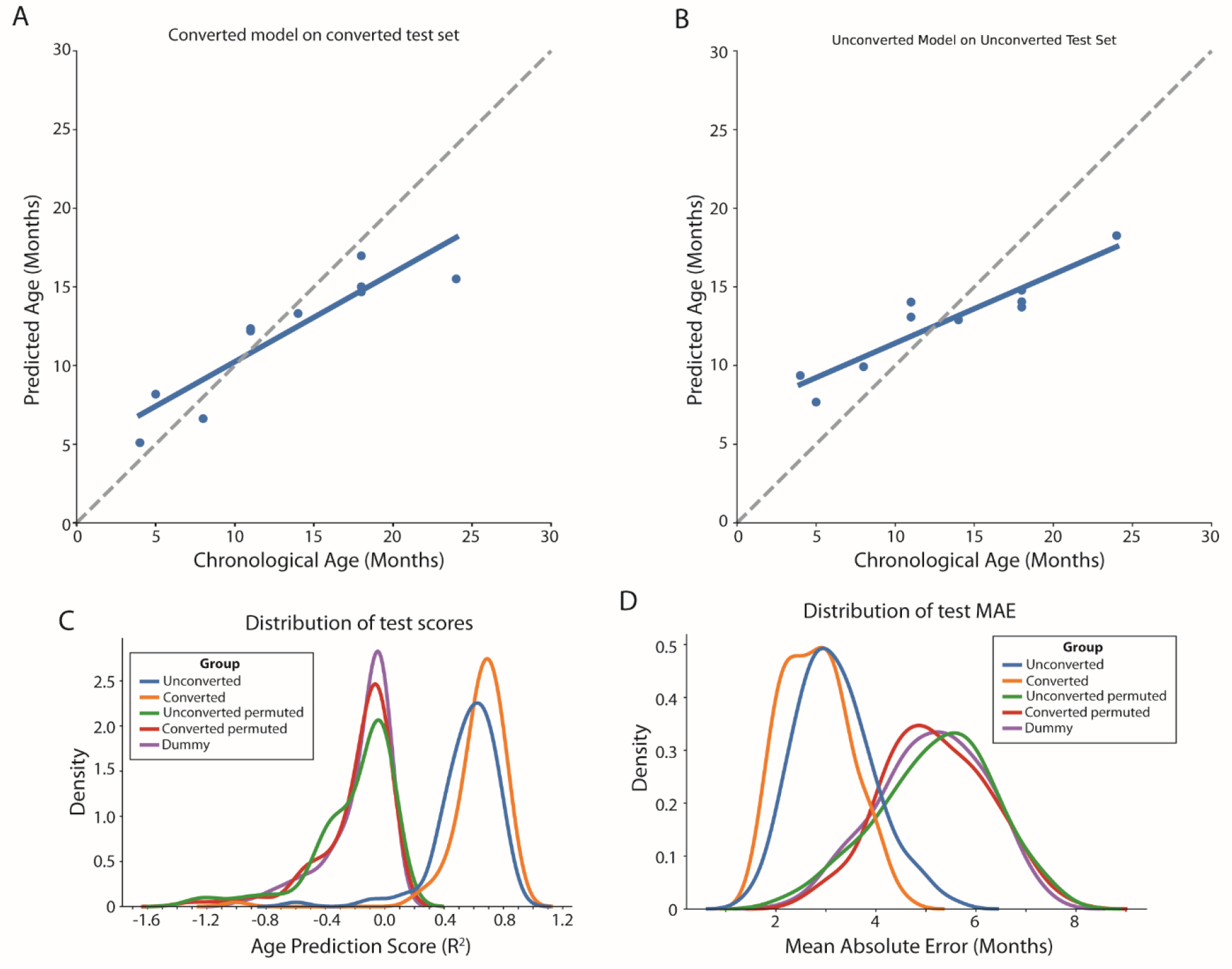
Converted and unconverted DNA can be applied to methylation arrays and generate equivalently predictive “epigenetic” clocks. Hippocampal DNA was isolated from dissected hippocampi of mice of various ages. In parallel, DNA was processed with standard bisulfite conversion (converted) or a mock conversion process lacking solely the bisulfite reagent (unconverted). Models were trained on both converted **(A)** and unconverted **(B)** data. This process was repeated for 100 iterations of generating new train/test splits and retraining to obtain distributions of prediction accuracy **(C)** and error **(D)**.

### Understanding the relationship between converted and unconverted epigenetic clocks

The observation that unconverted DNA could produce comparable epigenetic clocks led us to question if the clocks were detecting common signals shared between the converted and unconverted datasets, or if the underlying molecular changes were unique to each data type (Figure 4). To explore this, we first analyzed common clock sites of each data type with age (Figure 4A), as well as the correlation between array readouts and age from sites in each clock context (Figure 4B-D). We observed that most of the loci selected by one data type were not age predictive in the other. 13,138 loci passed our primary feature selection in converted samples, with 5,063 being selected by at least one clock iteration. Our unconverted dataset had 17,146 loci pass primary feature selection and 2,235 be used by one clock. When comparing the datasets, a small subset of loci common to both data types (49/556) were selected by even one of 100 clocks trained on either data type. Overall, the clocks selected fewer common loci than expected even by chance (hypergeometric p-value = 1.129 * 10^−21^). The correlations of the commonly selected loci with age were generally positive in the converted and negative in the unconverted (Figure 4C), but all combinations were observable (Suppl. Figure 1). The absolute differences in apparent “methylation” and their variability is noticeably larger in the unconverted samples (Figure 4A). These findings implicate unique, but not in dependent, molecular processes in generating age-predictive models from converted and unconverted DNA.

**Figure 4.**
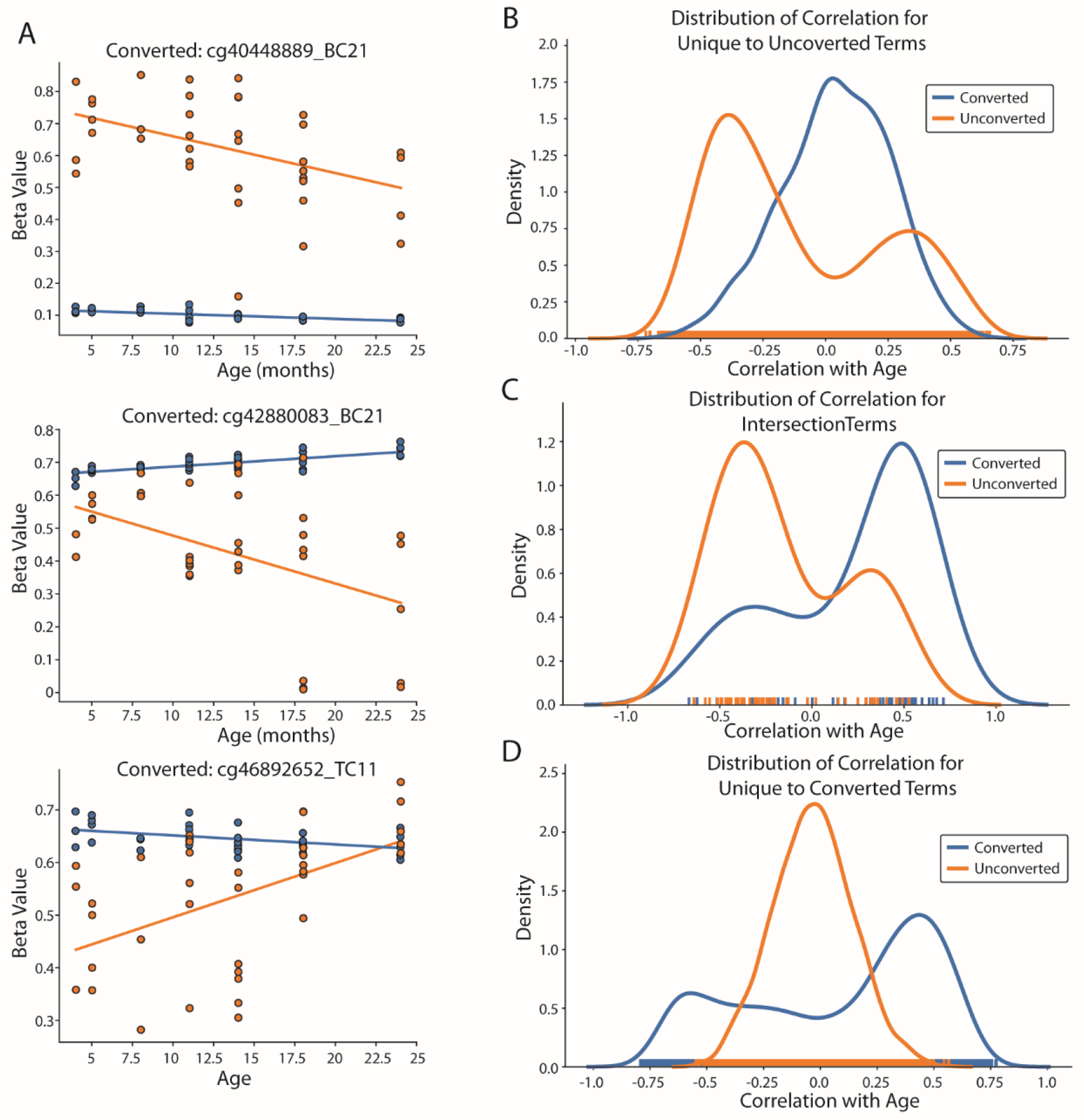
Converted and unconverted DNA are predominantly age-predictive due to unique signals. The loci selected by the previously described 100 clocks trained on converted and unconverted DNA were interrogated. A minority of sites (49 clock/556 primary selection) were common to only one clock in each data type. A majority in the converted (5,014/13,138) and unconverted (2,186/17,146) were unique. **A)** Three example loci that were age-predictive in both data types (**B - D)** distributions of the correlation between methylation from converted (blue) or “methylation” from unconverted (orange) and age. Distributions are shown for sites unique to unconverted clocks **(B)**, common to both **(C)**, or unique to converted **(D)**

To further explore this, we examined the distribution of univariate age associations in our primary feature selection (Figure 5A). This relationship also held in the loci selected by at least 1 clock in each data type (Figure 5B), where the probability of inclusion in the 100 model iterations was unrelated aside from one locus (cg47740109_TC21). We observed many loci with some age association (MI > 0.2) that were also unrelated to aging in the other data type (MI == 0). We asked how removing these loci with potentially confounded values would affect the epigenetic clock outputs. We hypothesized that removing the confounded loci would reduce epigenetic clock performance. Contrary to this, removing loci with non-zero age associations in the unconverted pairs increased epigenetic clock performance in terms of R^2^ (Figure 5C) and MAE (Figure 5D). Similarly, removing loci with methylation-derived age associations improved clocks trained on unconverted DNA. This implied that 1) clocks trained on either converted or unconverted DNA are age-predictive using unique biological signals, and 2) these signals are confounding each other’s age predictions in the correlated features.

**Figure 5.**
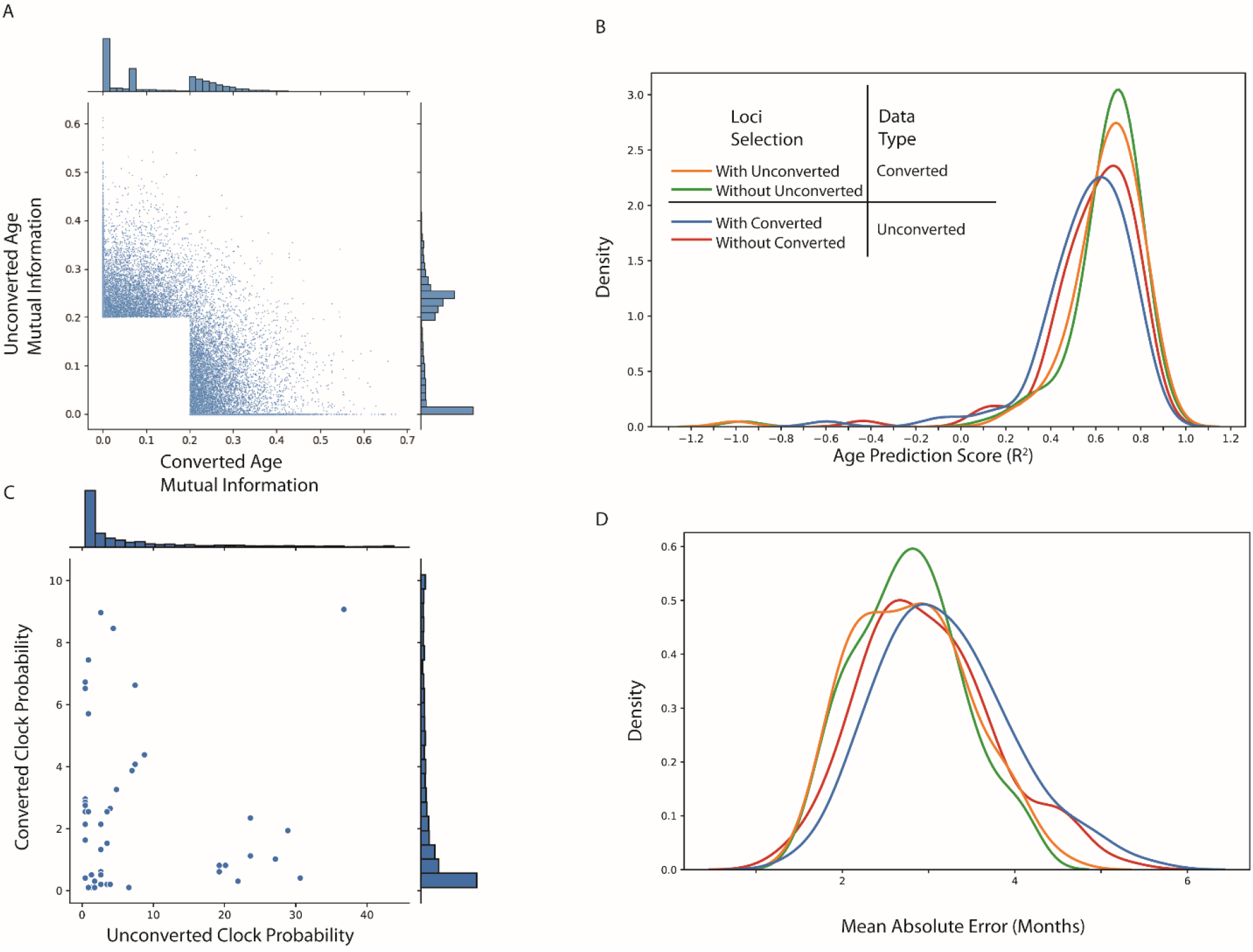
Isolating loci informative in bisulfite or unconverted samples improves clock performance for both data types. The loci selected by the previously described 100 clocks trained on converted and unconverted DNA were interrogated. **A** Joint mutual information distribution of sites passing primary feature selection from either converted or unconverted samples. **B** Probability of inclusion in a clock model for all common sites selected by both data types. **C and D** Filtering loci from the training data that contain any non-zero aging association from the opposite dataset (e.g. – loci with MI > 0.2 in converted and 0.0 in unconverted) does not significantly reduce model performance in terms of R^2^ (**C**) or MAE (**D**)

### III. Discussion

Our first set of experiments provided support for the hypothesis that somatic mutations can influence estimated DNA methylation levels in bisulfite-conversion assays. Effects of germline mutations have been previously described in the context of array data [18], but somatic mutations are more rarely studied alongside methylation – perhaps due to the unclear nature of their relationship [19]. In human sequencing data, it was found that methylated CpGs have fewer single nucleotide polymorphisms (SNPs) than unmethylated, but their relationship is more complicated than that binary change [29]. It was reported that loci from 20-60% methylation have higher SNP rates, which seems paradoxical given the nature of 5-methylcytosine’s direct deamination to thymine [14]. We expect that uracil-DNA glycosylases (UDGs) [30] would lead to increased rates of repair at cytosine > uracil mutations, but T:G mismatches could be more error-prone. Enzymes such as thymine DNA glycosylase (TDG) and methyl binding domain 4 (MBD4) repair the T:G mismatches caused by mC=>T mutations [31]. However, there is a conserved reduction in thymine excision activity compared to uracil excision activity at T:G and U:G mismatches, respectively. The conserved impairment of glycosylase activity is thought to protect against aberrant base excision at A:T base pairs which are abundant throughout the genome, whereas dU is rare and typically a byproduct of mutagenic DNA damage. Therefore, the activity of these enzymes does not fully protect against C=>T mutations, as evidenced by the prevalence of C=>T mutations in cancer[32] and aging[33]. The relationship between somatic mutations and age have been explored in the cancer field[34], and the relationship is also dependent on TP53 mutations [35]. The idea that these mutations affect methylation readouts is also discussed [36, 37], but the direct evidence that the mutation itself is read out as loss of methylation has not been tested. This has potentially interesting ramifications in epigenetic clock studies, as the potential for mutations to influence methylation data means that clocks may contain information from both mutations and methylation which are currently undetermined.

Improving our understanding of the mechanisms behind epigenetic clock formation will depend upon our ability to disentangle methylation changes from somatic mutations in the aging context. We see two ways forward in this regard: specially designed short-read sequencing approaches like double-stranded bisulfite sequencing [38] or five-letter seq[39] will enable us to isolate these two variables by assaying base pairs in a single DNA molecule instead of relying on the reference genome alone. This approach, after bisulfite conversion, will lead to T:G pairs at unmodified cytosines, and A:T pairs at mutated bases.

Another approach is removing the need for bisulfite treatment and post-conversion treatments entirely, which have complex, and perhaps some uncharacterized, effects on DNA bases. Nanopore sequencing could be leveraged to isolate mutations, methylation, and other DNA modifications and by-products of DNA damage simultaneously [40]. There might also be many other modifications present in real DNA samples with aging which influence bisulfite-based assays. Regardless of approach, it is obvious that bisulfite methods make assumptions that are likely violated in cancer, and the field is actively searching for alternative solutions. The relationship between these same processes and aging will be of great importance to understanding epigenetic clocks and to separate the unmodifiable somatic mutations from therapeutically tractable DNA methylation alterations.

Our next set of experiments was targeted at understanding how signals other than methylation may influence the ability of methylation array data to predict chronological age. The approach of mock conversion allows us to capture any variability that is introduced by the other components of the bisulfite preparation but does not allow us to directly measure mutations in general, nor C>T mutations. The data from sequencing reflects the expected results of bisulfite assumptions (Figure 1), but the array data presents another distribution entirely (Supplemental figure 2). Nonetheless, these results provide evidence that epigenetic data suffers from being unable to distinguish between real methylation changes and other phenomena that alter the microarray signal. An obvious example is found in the delineation of mC from hydroxymethylcytosine (hmC), where we have previously described how disentangling cytosine modifications can reveal physiological changes otherwise masked by bisulfite preparations [41]. Our results also imply that the relationship between our unconverted library predictions and standard methylation data is unlikely to be due solely to mutations. However, we were incorrect in our initial hypothesis that controlling for the unconverted signal would reduce clock performance (Figure 5). Our findings provide evidence for both a path to adjusting epigenetic clocks to delineate real methylation changes with age, as well as evidence that some other genomic modification provides an alternative aging biomarker with comparable prediction accuracy.

In conclusion, bisulfite studies in sequencing and array contexts are confounded by molecular changes other than DNA methylation alone. In the sequencing context, we demonstrate a clear relationship between mutations and methylation changes, which are likely relevant in both age prediction and DNA methylation research in general. Meanwhile, we show that arrays do not need bisulfite conversion to produce age-predictive signals. These results provide direction for adjusting future clock studies, as well as focusing epigenetic theories of aging on true epigenetic alterations and avoiding the litany of confounding variables that are currently ignored.

## IV. Methods

### Cancer Cell Culture

Immortalized young adult mouse colon (YAMC) epithelial cells were assigned to control or treatment groups. Treated cells were transformed as previously described [42]. Briefly, cells were co-cultured with macrophages polarized by Enterococcus faecalis infection or exposed to trans-4-hydroxy-2-nonenal (4-HNE). Macrophage exposure occurs in a dual-chamber system for 72 hours with 96 hours of recovery, while 1 µM 4-HNE exposure is done in one hour treatment windows with 1 week of recovery repeated over 16 treatments.

### Mouse Sample Collection

C57BL/6 mice were obtained from the National Institute on Aging (NIA) aging colony at 4-18 months of age. Some mice were aged additionally for collection, up to 24 months of age. Mice were sacrificed at 4, 5, 8, 11, 14, 18, and 24 months of age, then hippocampi were harvested, snap frozen in liquid nitrogen, and stored at –80 °C until used. All animal experiments were performed according to protocols approved by the OMRF Institutional Animal Care and Use Committee.

### Whole-Genome Shotgun Sequencing

Whole genomic shotgun (WGS) libraries were prepared using Swift Biosciences Turbo v2 library preparation kits as per the manufacturers protocol (Swift Biosciences, Ann Arbor, MI). Briefly, DNA from each respective sample was simultaneously fragmented, end-repaired, and treated to create a single A-base overhang on each end. Following an SPRI bead-based cleanup the DNA fragments were ligated to a unique-dual-indexed adapter set. The prepared libraries were fluorometrically quantified with a Qubit fluorometer (Life Technologies, Grand Island, NY) and pooled at equimolar concentrations. The pool was then precisely quantified via qPCR with a NEBNext Library Quant Kit for Illumina (New England Biolabs, Ipswich, MA) on a Lightcycler 480 instrument (Roche Diagnostics, Indianapolis, IN). The pool was then sequenced on an Illumina NovaSeq 6000 instrument using an S4 flow cell and paired-end 150bp reads.

### Bisulfite-Capture Sequencing

Bisulfute Oligonucleotide Capture Sequencing (BOCS) was performed utilizing genomic DNA which is made into a sequencing library followed by bisulfite conversion. After conversion the target genomic regions of interest are captured with probes against methylated and unmethylated versions of the targeted genomic regions. The target regions are then captured and amplified prior to sequencing. Targeted regions of the mouse genome include 109Mb containing most of the annotated promoters and CpG islands. In total nearly 3 million CG and 28 million CH sites are in the targeted regions.

### Methylation Microarrays

Isolated hippocampal DNA was assigned to one of three groups: bisulfite-converted (Converted), mock bisulfite treatment (Unconverted. Bisulfite-converted samples were treated using the Zymo EZ DNA Methylation Kit following the manufacturer protocol with minor modifications by Illumina [23]. Unconverted samples were treated identically without addition of bisulfite reagent. Sample DNA was then enzymatically fragmented and hybridized to the BeadChip and washed before extension, staining, and imaging. Imaging-generated .idat files were then parsed into matrices of methylation loci X sample using methylprep [43]. Missing values were imputed using sklearn’s KNN Imputer [44].

### Epigenetic Clock Training and Analysis

Epigenetic clocks were fit as previously described [25] using sklearn’s elastic net regression implementation with data split into 75% test and 25% train sets. For repeated clock trainings, the random state and train/test split were altered between each replicate, with 100 models being fit for each data type and data subset. Training was done using 2-fold cross validation and maxiter = 5000. Primary feature selection was done using sklearn’s mutual information regression with the target of chronological age in months, excluding sites with computed MI <0.20. Over/underrepresentation of loci in clock models was computed using the hypergeometic test implemented in scipy stats [45].

### Pseudomethylation Analysis

WGSS fastq’s were aligned to the mm10 reference genome using Bowtie 2 [46] or Bismark[27] with --sensitive parameters (N = 1, L = 20). Bismark’s methylation extractor was used to make methylation calls from both bisulfite sequencing data (deamination by bisulfite) and whole genome sequencing data (deamination by mutation). Mutated regions were determined using Bowtie 2 samples aligned to the mm10 reference genome with –sensitive parameters. Variants were called using samtools mpileup and bcftools call.

### Use of AI

ChatGPT was used to help refactor code for generating publication-quality figures. Code was primarily used for generating plots or reshaping data from one format to another for plotting. Most prompts consisted of pre-existing code with the request of refactoring into functions or reformatting data consistently across datasets.

## Acknowledgements

This work was funded by NIH grants #P30AG050911 (JDW, WMF), #P30GM149376 (JDW), and #R01-CA230641 (MMH). This project was also supported by a Longevity Impetus Grant to JDW and WMF.

## Author Contributions

Study Design – HLP, WMF, JDW, WH. Wet-lab experimentation: VAA, RBU. Statistical analysis and machine learning – HLP, WH. Programming support – CBG, CAB, DS. Manuscript Preparation – HLP, WH, JDW, WMF. Supervision and funding – JDW, WMF, HLP, and MH.

## Supplementary Figures

**Supplementary Figure 1.**
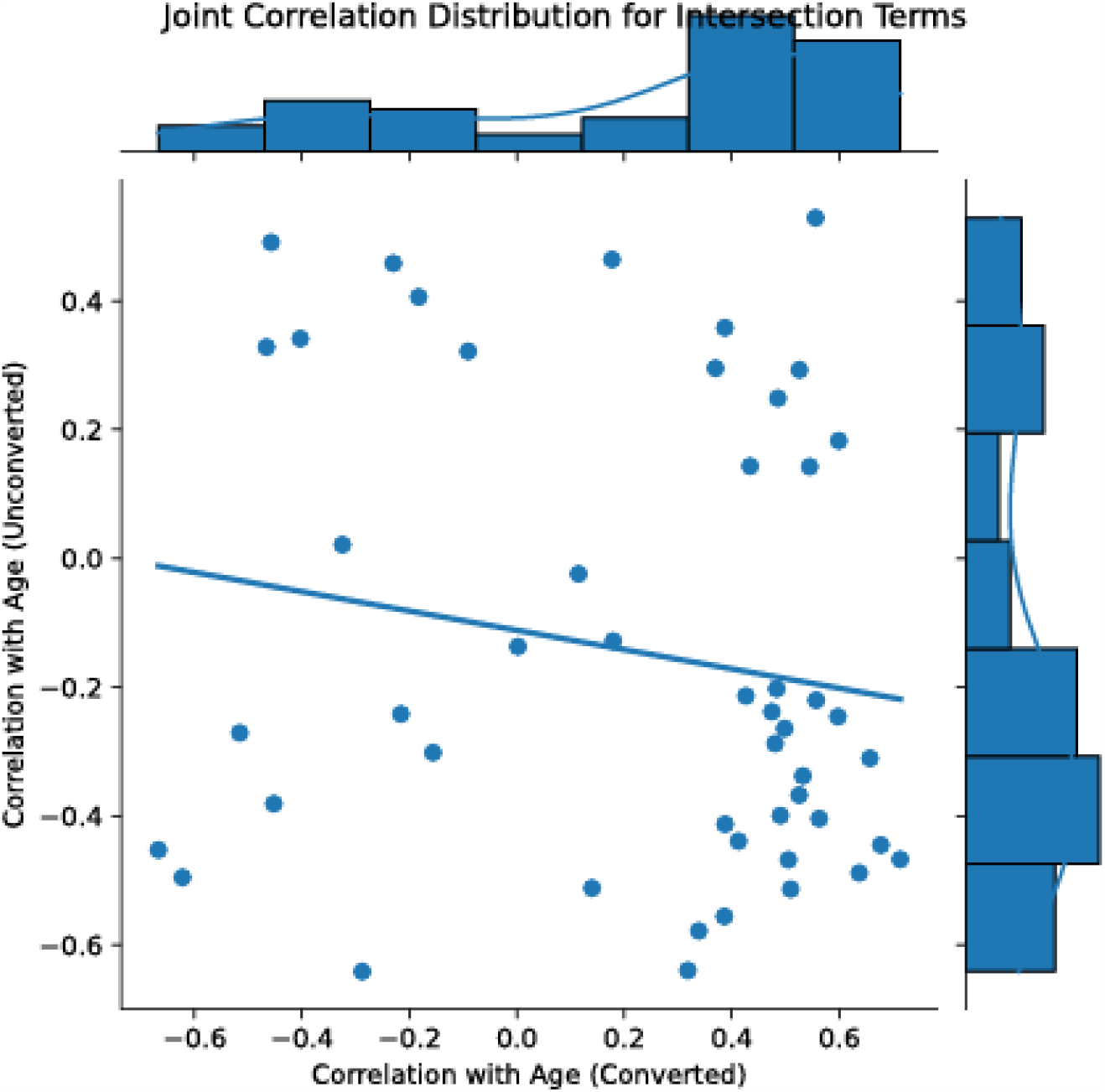
Jointplot of age correlations of individual loci that were common to at least 1 unconverted and converted clock

**Supplementary Figure 2.**
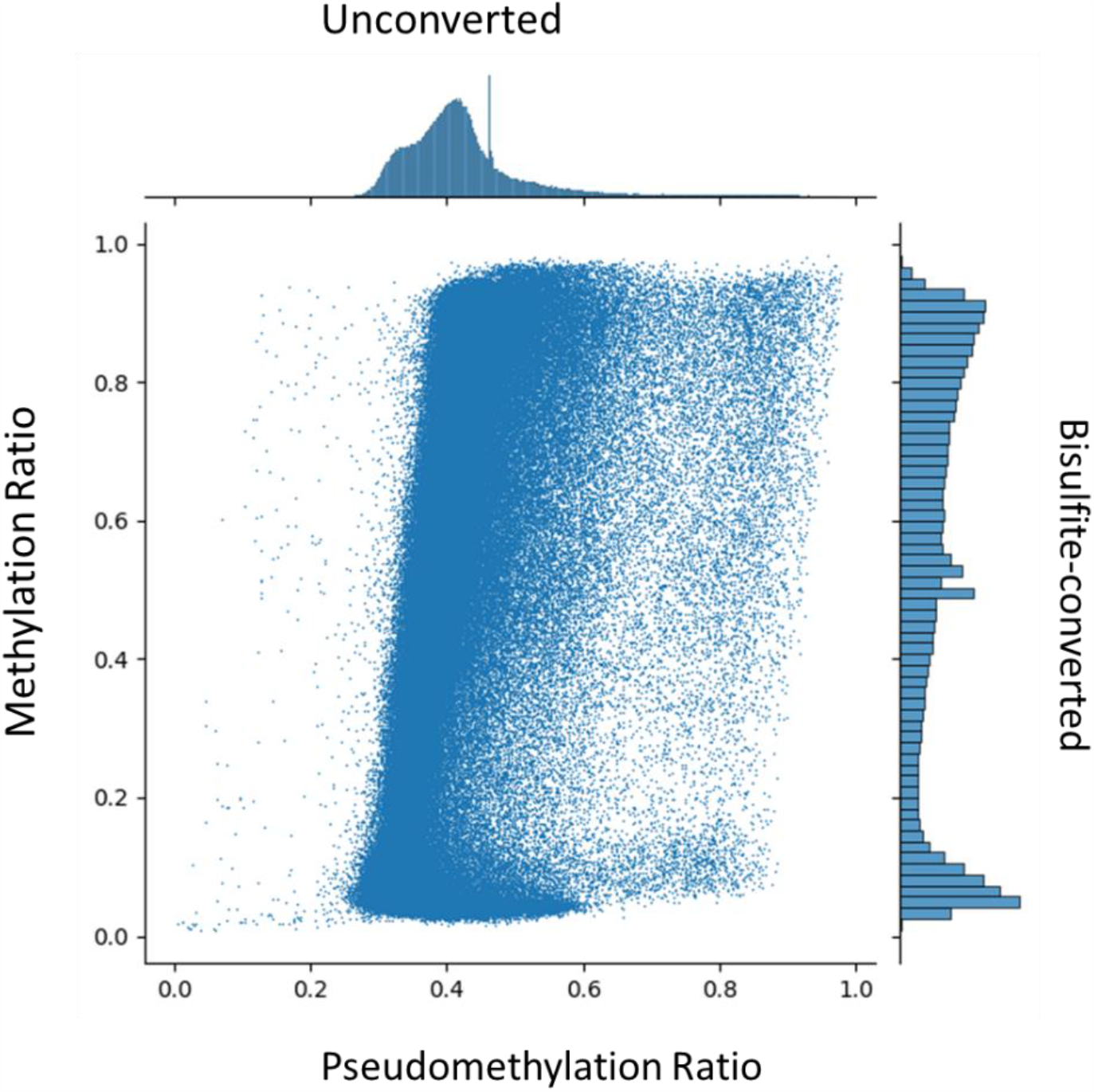
Comparison of converted and unconverted signal from mouse methylation arrays after preprocessing

